# Computational modeling of dorsal root ganglion stimulation using an Injectrode

**DOI:** 10.1101/2023.09.20.558675

**Authors:** Sauradeep Bhowmick, Robert D. Graham, Nishant Verma, James K. Trevathan, Manfred Franke, Stephan Nieuwoudt, Lee E. Fisher, Andrew J. Shoffstall, Douglas J. Weber, Kip A. Ludwig, Scott F. Lempka

## Abstract

**Objective:** Minimally invasive neuromodulation therapies like the Injectrode, which is composed of a tightly wound polymer-coated platinum/iridium microcoil, offer a low-risk approach for administering electrical stimulation to the dorsal root ganglion (DRG). This flexible electrode is aimed to conform to the DRG. The stimulation occurs through a transcutaneous electrical stimulation (TES) patch, which subsequently transmits the stimulation to the Injectrode via a subcutaneous metal collector. However, effectiveness of stimulation relies on the specific geometrical configurations of the Injectrode-collector-patch system. Hence, there is a need to investigate which design parameters influence the activation of targeted neural structures.

**Approach:** We employed a hybrid computational modeling approach to analyze the impact of the Injectrode system design parameters on charge delivery and the neural response to stimulation. We constructed multiple finite element method models of DRG stimulation and multi-compartment models of DRG neurons. We simulated the neural responses using parameters based on prior acute preclinical experiments. Additionally, we developed multiple human-scale computational models of DRG stimulation to investigate how design parameters like Injectrode size and orientation influenced neural activation thresholds.

**Main results:** Our findings were in accordance with acute experimental measurements and indicated that the Injectrode system predominantly engages large-diameter afferents (Aβ-fibers). These activation thresholds were contingent upon the surface area of the Injectrode. As the charge density decreased due to increasing surface area, there was a corresponding expansion in the stimulation amplitude range before triggering any pain-related mechanoreceptor (Aδ-fibers) activity.

**Significance:** The Injectrode demonstrates potential as a viable technology for minimally invasive stimulation of the DRG. Our findings indicate that utilizing a larger surface area Injectrode enhances the therapeutic margin, effectively distinguishing the desired Aβ activation from the undesired Aδ-fiber activation.

## INTRODUCTION

Neuromodulation devices, such as spinal cord and dorsal root ganglia stimulators hold great potential for treating debilitating chronic pain, which is one of the largest public health challenges in the United States, affecting over 100 million Americans and accounting for more than $600 billion in healthcare cost and lost productivity [1,2]. The goal of neurostimulation therapies is to apply exogenous electric fields to the nervous system to elicit a desired therapeutic response and improve quality of life. Conventional pain management approaches, including the use of opioids, have unfortunately contributed to a concerning rise in addiction and subsequent fatal overdoses. Recent data shows a doubling in overdose cases, underscoring the urgent necessity for alternative and non-addictive pain therapies [3,4,5].

Spinal cord stimulation (SCS) has been a widely used neurostimulation therapy for treating intractable neuropathic pain in the trunk and limbs [2]. SCS involves implantation of one or more electrode arrays in the spinal epidural space and applying brief electrical impulses to create analgesia, putatively through pain-gating mechanisms within the spinal cord [6, 7]. Despite the widespread success of SCS in treating several chronic pain conditions, pain in specific areas, such as the groin, foot, low back, and knee, can be difficult to target due to the complex anatomy of the spinal cord. Several other factors, such as posture-related motion of the spinal cord in the thecal sac, lead migration, and electrical shunting in the cerebrospinal fluid (CSF), can limit successful neural targeting with SCS [3,8,9]. Therefore, for patients with pain in regions that are difficult to target with SCS, dorsal root ganglion stimulation (DRGS) can be considered as a viable alternative [3,4,5].

The dorsal root ganglion (DRG) is located near or within the foramen in the posterior spinal root at each level of the spinal cord. Each DRG contains the cell bodies of all the primary sensory neurons, and a portion of the axons innervating a single dermatome [10]. DRG neurons are pseudounipolar: a single axon process extends from the soma, bifurcates at a large node of Ranvier called the T-junction, and forms an axon that projects to the spinal cord and an axon that extends to the periphery [11]. Due to the precise targeting of a single dermatome’s primary afferents, DRGS can provide patients with focal, dermatome-specific pain relief. In contrast to SCS electrodes which are placed along the dorsal aspect of the spinal cord, DRGS involves implantation of the annular electrode arrays in the intraforaminal space next to the DRG. DRGS was approved by the US Food and Drug Administration in 2016 to treat intractable complex regional pain syndrome (CRPS) of the lower limbs [14] and has been subsequently used for several other pain etiologies (e.g., phantom limb pain, painful diabetic neuropathy, groin pain) [15–18]. Early reports showed that due to the compactness of the intraforaminal space and scarcity of CSF around the ganglion [12], DRGS electrode arrays may be less prone to the type of lead migration and postural effects that can decrease the efficacy of SCS [13].

Both SCS and DRGS are limited by the procedures required to implant leads in epidural space. This process can be uncomfortable for patients, and permanent implants often require the use of general anesthesia. Furthermore, over a period of 4 years, the FDA has reported a total of 107,728 medical device reports related to SCS for pain, including 497 associated with a patient death, 77,937 with patient injury, and 29,294 with device malfunction [52]. Additionally, recent clinical reports have shown significant DRG lead migration at the sacral level when using a transforaminal approach [19–20], and several studies have reported other complications, such as pain near the implantable pulse generator and lead fracture [21–23], resulting in some countries pausing DRGS implantations. Thus, to achieve improved clinical implementation and to reduce complexity to minimize the failure points, a minimally invasive procedure is needed to deliver effective stimulation without the drawbacks of existing electrode technologies and their placement procedures.

The Injectrode is a tightly wound polymer coated platinum/iridium microcoil which is injected via an 18-gauge needle near a neural target where it forms a highly conforming, flexible, clinical-grade electrode platform [26,54]. The Injectrode consists of three continuous regions: an uninsulated portion at one end to serve as a stimulation site, an insulated lead portion in the middle, and another uninsulated portion at the other end to serve as a subcutaneous charge collector [43]. During delivery, the Injectrode can fold into a variety of conformations dependent on the target anatomy. The delivered electrode is therefore larger in diameter than the needle from which it was deployed, thereby reducing the chances of migration. Once the Injectrode is delivered to the target structure, then the insulated lead is extruded back to the superficial tissue layers where the subcutaneous charge collector is placed [43,53]. Electrical current is delivered transcutaneously to the charge collector located directly beneath the skin surface using noninvasive skin adhesive patch electrodes, eliminating the need for implantable pulse generators or wires perforating the skin. From the charge collectors, the electrical currents then travel through the insulated lead wire to the Injectrode situated on top of and near the DRG in the intraforaminal space. Thus, it is important to understand the charge delivery pathway in this electrical stimulation system because it directly affects the safety and effectiveness of the therapy.

To gain a comprehensive understanding of the charge delivery mechanisms employed by this system, it is important to first examine the impact of various technical and anatomical factors on the electrical stimulation delivered to the DRG. These factors include the size of the Injectrode and the relative placement and orientation of the patch electrodes and the collector. Although similar studies have been conducted on preclinical models of vagus nerve stimulation using the Injectrode [43], due to the substantial differences in geometry and anatomical positioning between the DRG and the vagus nerve, along with their respective surrounding soft tissues, the associated side effects exhibit considerable variation. When targeting the DRG, there is a potential risk of stimulating sensory Aδ-fibers, which can induce acute pain sensations. Thus, the effect of clinically adjustable parameters on neural recruitment during DRGS with the Injectrode system remains unclear. Therefore, leveraging computational modeling to bridge this knowledge gap could optimize the clinical efficacy of the system and enable the exploration of different configurations.

In this work, we implemented a computational modeling approach to investigate the effects of clinically relevant factors on neural recruitment during DRGS via the Injectrode. Our initial goal was to validate our computational modeling approach by comparing experimental measurements to the corresponding model predictions. Therefore, we first developed a computational model of DRGS in a feline model, compared and validated our model predictions to the neural recruitment observed in our previous experimental work [24]. After validating our computational modeling approach, we then developed a generalized model of stimulation of the human L5 DRG, a common stimulation target to manage chronic foot pain [16–17,25]. We used this clinical-scale model to examine how different shapes, sizes, and orientations of the Injectrode affected neural recruitment profiles within the DRG. Finally, we built a full body human model of bilateral DRGS with a complete bipolar Injectrode system, i.e., skin electrodes, collectors, microwires, and Injectrodes. It was observed that for full body bipolar TES-collector-Injectrode stimulation, the activation thresholds were significantly lower for an Injectrode of larger surface area suggesting that placing the Injectrode such that it covers a maximal area possible may be the optimal configuration for activating relevant neural tissue because it led to an amplified therapeutic window differentiating the desired Aβ-fiber activation from undesired Aδ-fiber activation. In contrast, smaller Injectrode sizes exhibited characteristics more reminiscent of conventional SCS/DRG electrodes, resembling a point-source effect.

## MATERIALS AND METHODS

We developed computational models of DRGS to investigate how the model predictions compared to experimental data and how clinically controllable factors (e.g., Injectrode size) affect neural activation in the DRG. Since DRGS is believed to provide analgesia via pain-gating mechanisms induced by the activation of Aβ-fibers [3–5], we built computational models to study neural recruitment of Aβ-fibers within the DRG using the Injectrode. For the canonical and the full-body human-scale models, we also built computational models of Aδ-fibers to examine possible off-target effects induced by Injectrode stimulation. We coupled a finite element method (FEM) model of a lumbar DRG to multicompartment models of sensory Aβ- and Aδ-fibers [5]. We used the FEM model to calculate the potential fields generated by DRGS and applied these potentials to the multicompartment neuron models (Figure 1). We examined the stimulation amplitudes required to activate Aβ- and Aδ-fibers placed throughout the dorsal aspect of the DRG. We also investigated how the neural recruitment profiles changed as a function of stimulus pulse width and Injectrode geometry.

**Figure 1:**
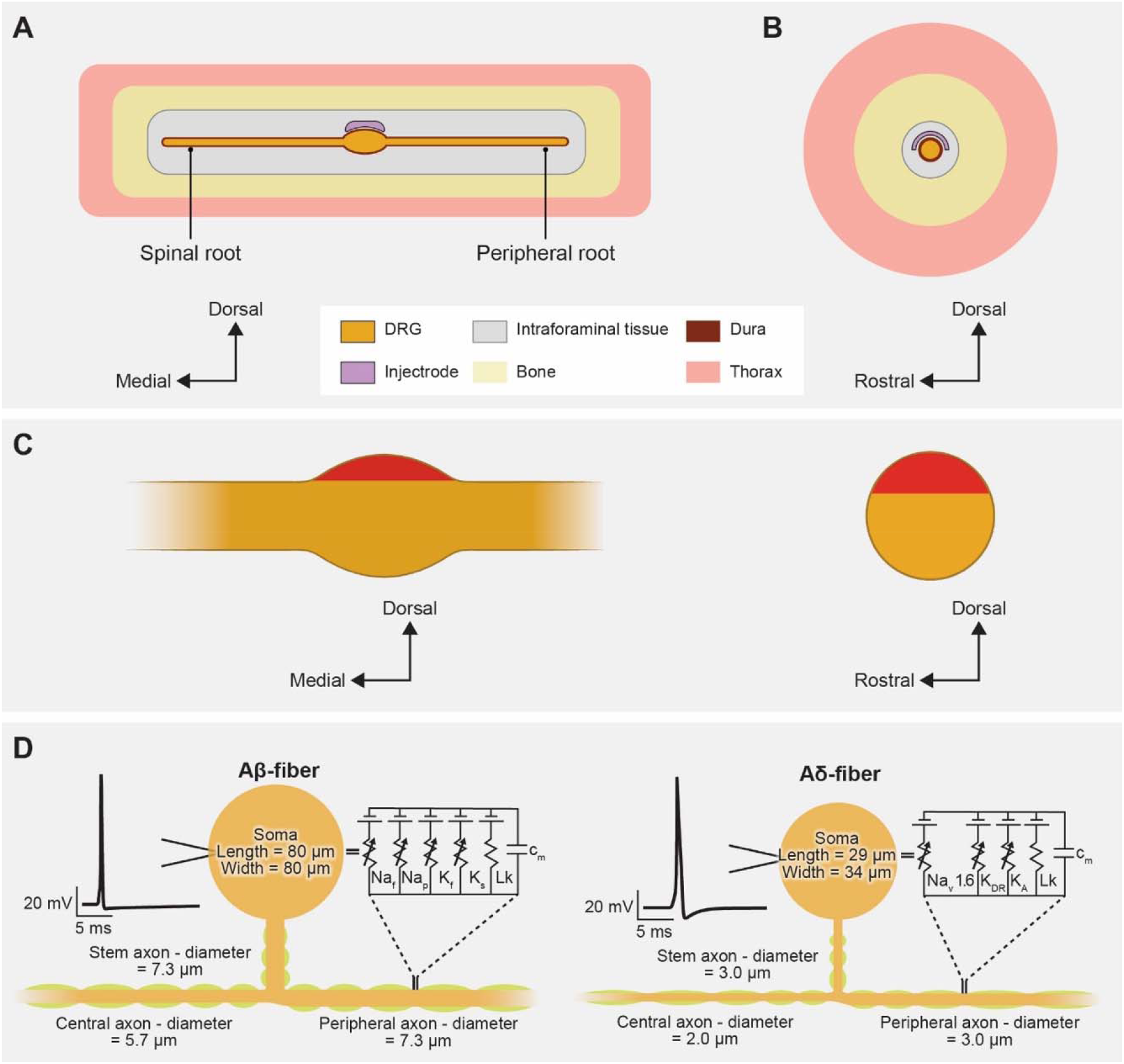
Representative schematic of our finite element method (FEM) model of DRGS. We developed an FEM model of a DRG and surrounding anatomy. Two separate versions of this model were developed and scaled to fit dimensions of the feline L7 DRG and human L5 DRG. (A) Side view of the DRG with the Injectrode oriented above the ganglion. (B) Cross-sectional view through the middle of the DRG and Injectrode. (C) Red-shaded regions indicate the locations of the somata of primary sensory neurons in sagittal and transverse DRG cross sections. (D) Multicompartment models of primary sensory neurons, representing the pseudounipolar morphology of a large-diameter myelinated Aβ-fiber and a smaller-diameter thinly myelinated Aδ-fiber. An example action potential from each model neuron is shown on the left. The equivalent circuit diagram with active ion conductances included in the nodal, initial segment, and soma compartments is shown on the right.

We constructed three-dimensional FEM models using anatomical data from existing literature. We used previous computational modeling studies and experimental measurements [28–31] to assign electrical conductivities for each tissue type. We modeled each tissue as having an isotropic conductivity, except the nerve root, which we modeled as having an anisotropic conductivity (Table 1).

**Table 1.** Electrical Conductivities assigned to the anatomical components in the FEM models.

| Parameter | Value (S/m) | Reference |
| --- | --- | --- |
| Gray matter | 0.230 | [29] |
| White matter (Longitudinal) | 0.600 | [29] |
| White matter (Transverse) | 0.083 | [29] |
| Dural covering | 0.600 | [31] |
| Bone | 0.020 | [28] |
| General tissue | 0.250 | [29] |
| Fat | 0.04 | [44] |
| Encapsulation | 0.170 | [30] |
| Skin | 0.148 | [47] |
| Platinum | $9.43 \times 10^6$ | [48] |

To calculate the potential fields generated by DRGS, we used COMSOL Multiphysics (COMSOL, Inc., USA) and applied the relevant Dirichlet and Neumann boundary conditions. To simulate a canonical DRG model of monopolar configuration, we applied a unit current stimulation boundary condition (i.e., 1 A) to the surface of the Injectrode and set the outer boundaries of the general thorax domain to ground (i.e., 0V). We used the conjugate gradient method to solve the Laplace equation:

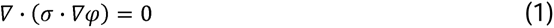

where σ is the tissue stiffness matrix and φ is the calculated electric potential.

We stimulated multicompartment models of primary sensory neurons found in the DRG using the NEURON simulation environment (v7.4) (32-33). We implemented previously published models of an Aβ-fiber (Figure 1) [4–5]. Following our previous work [4,5], we set our Aβ-fiber model central axon diameter to 5.7 μm and the peripheral axon diameter to 7.3 μm [35]. The models had a soma 80 μm long and 80 μm wide, connected to a 7.3-μm diameter stem axon. The stem axon extended 789 μm before splitting into two axons. The myelinated compartments consisted of two concentric layers containing linear leak conductances with a parallel membrane capacitance. The nodes of Ranvier contained the parallel active nodal conductances of the sensory-specific axons described by [36]: fast Na^+^, persistent Na^+^, fast K^+^, and slow K^+^ ion channels. The active nodal conductances were modeled in parallel with a linear leakage conductance and membrane capacitance (Figure 1D). In line with our previous work, the channel densities at the somata were lower compared to the nodes [4–5]. For the canonical and full body human models, we implemented the low-threshold mechanoreceptor (LTMR) Aδ-fiber models that were also previously developed in computational modeling work from our group [5]. The thinly myelinated medium-diameter Aδ-fibers express distinct voltage-gated sodium channel profiles, namely Nav1.6, similar to other non-nociceptive myelinated mechanoreceptors [5]. Each Aδ-fiber model had a soma 29 μm long and 34 μm wide, connected to a 3.0-μm diameter stem axon. The stem axon extended 840 μm before splitting into two axons. One axon projected towards the spinal cord, with a diameter of 2.0 μm, while the other projected to the periphery and had the same diameter as the stem axon (i.e., 3.0 μm) (Figure 1D).

We linearly interpolated the extracellular potentials calculated in Equation (1) onto the middle of each compartment of the cell models. We applied the extracellular potentials to the multicompartment models using NEURON’s extracellular mechanism within the Python programming language [33]. We calculated each compartment’s time-dependent membrane voltage in response to DRGS by using a backward Euler implicit integration method with a time step of 5 μs (Figure 2). The tissue conductivities of the FEM model were linear. Therefore, the potential field generated by a specific DRGS amplitude was a scalar multiple of the potential field generated by a unit (i.e., 1 A) stimulus. We calculated activation thresholds for anodic/cathodic pulses using a binary search algorithm with a resolution of 0.1 μA [4–5].

**Figure 2:**
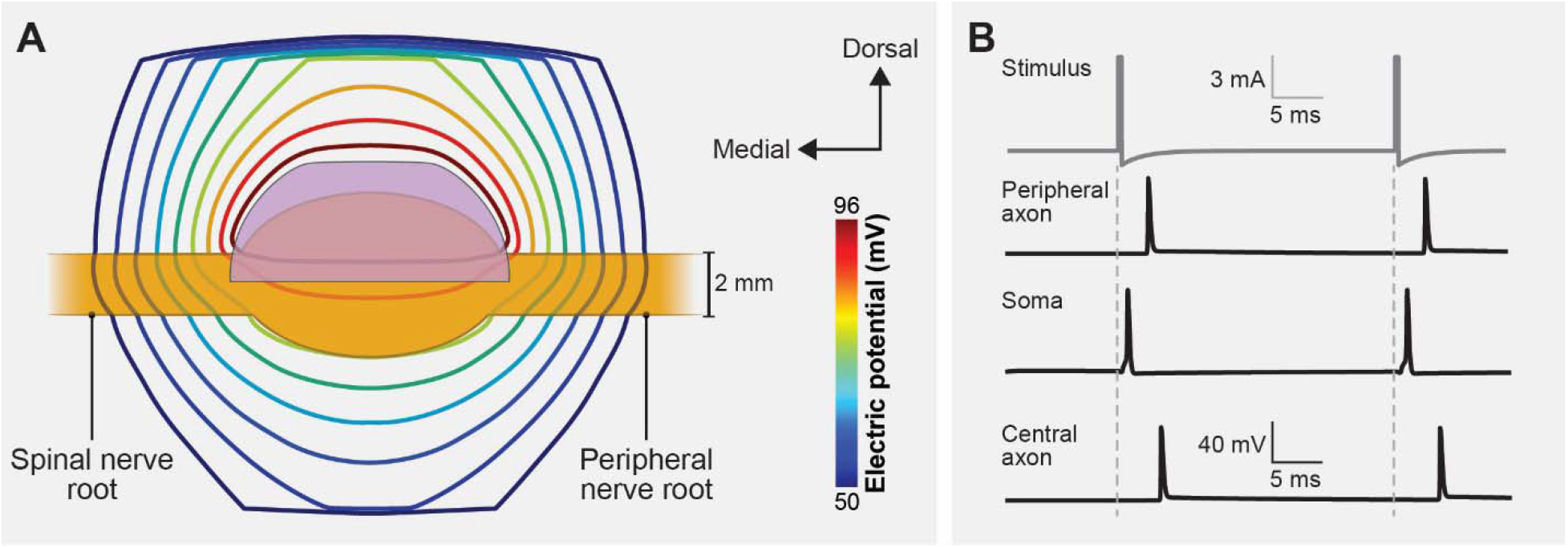
Coupling the finite element method (FEM) model of a human L5 DRG to the multicompartment models of primary sensory neurons. (A) Isopotential lines of the extracellular potentials generated by monopolar DRGS calculated from the FEM model. (B) Time-dependent transmembrane voltages resulting from stimulating an example Aβ-fiber with a stimulus having a pulse amplitude of 6 mA, pulse width of 300 µs, and pulse frequency of 40 Hz.

### Feline model of DRGS

To validate our computational modeling approach, we developed a canonical computational model of monopolar stimulation of a feline L7 DRG with the Injectrode and subsequently compared our findings with a set of experimental results [24]. We constructed a three-dimensional FEM model (Figure 1) based on measurements from previous computational modeling studies that utilized cadaver and imaging studies of the DRG and surrounding anatomy (e.g., dural covering, intraforaminal tissue, bone) (Table 2) [4–5, 27]. We built this FEM model in the 3-matic module within the Mimic Innovations Suite (Materialise, Belgium). We modeled the Injectrode to have a surface area of 48 mm^2^ that replicated the average size used in our previous experimental work [24]. We imported the volume mesh generated in 3-matic into COMSOL Multiphysics and applied 1 A at the Injectrode surface, grounded the outer surfaces of the model, and solved Equation (1). We interpolated the model solutions into the center of each compartment of the multicompartment neuron models. We validated the model by comparing the minimum stimulation current required to invoke a response in the DRG neurons to the current needed to produce a measurable electroneurogram response in the preclinical experiments [24]. For each set of stimulation parameters, we calculated the minimum stimulus amplitude necessary to elicit one or more action potentials in each Aβ-fiber (i.e., the activation threshold). We populated the dorsal aspect of the feline L7 DRG with 1355 Aβ-fibers spaced 200 µm apart, with the somata lying near the dorsal surface (Figure 3A). To mimic the experimental conditions, we used symmetric biphasic stimulus pulses applied at a frequency of 58 Hz and three different pulse widths of 80, 150, and 300 μs [24].

**Figure 3:**
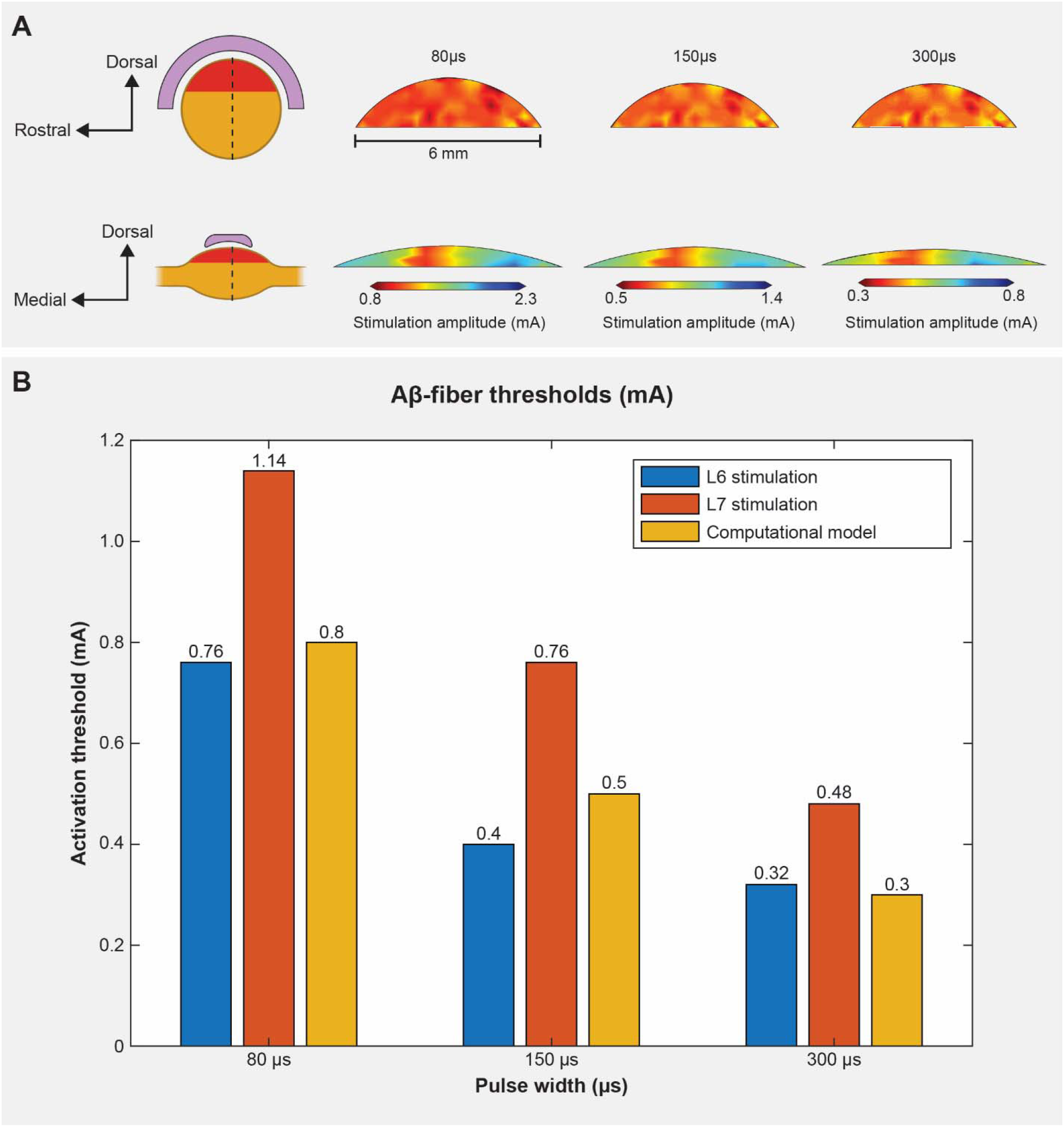
DRGS amplitudes required to elicit one or more action potentials (activation threshold) in Aβ-fibers for stimulation with an Injectrode in the feline model. (A) The contour plots show activation thresholds along the dorsal-rostral plane and the dorsal-medial plane for three different pulse widths. The red shaded region indicates the location of the somata of the primary sensory neurons enclosed by the Injectrode at the top. (B) Comparison of activation thresholds generated by our computational model with the ECAP thresholds from the acute experiments (for two lumbar levels) across three different pulse widths [24].

**Table 2.** Dimensions of the canonical FEM model of the feline L7 DRG.

| Parameter | Value | Reference |
| --- | --- | --- |
| DRG length | 6.50 mm | [27] |
| DRG width | 3.05 mm | [27] |
| Nerve root radius | 0.75 mm | [27] |
| Dural sheath thickness | 0.02 mm | [27] |

### Canonical human model of DRGS

To examine how the Injectrode geometry affects neural recruitment profiles during DRGS and assess its potential adverse effects associated with the activation of acute pain fibers, we developed a canonical model of DRGS applied to a human L5 DRG (Figure 4A). We based the model geometry on our prior work (Table 3) [4–5], with the standard clinical annular DRGS electrode array replaced by the Injectrode. The FEM model schematic in Figure 1 was scaled to represent the dimensions of a human L5 DRG (Figure 4A). We varied the included angle covered by the Injectrode on the dorsal-rostral plane (φ) and dorsal-medial plane (θ) from 30 degrees to 150 degrees at an interval of 60 degrees, thereby generating a total of nine different Injectrode geometries (Table 4; Figure 4). The Injectrodes were embedded in a 300 μm thick encapsulation layer, to represent typical foreign body response to implanted materials [30]. As described in the previous sections, we developed an FEM model to mimic monopolar stimulation conditions and interpolated the model potential fields onto the center of each compartment of the Aβ- and Aδ-fibers within the human L5 DRG model. We uniformly populated the dorsal aspect of the L5 DRG with 1378 Aβ-fibers and 1378 Aδ-fibers spaced 300 µm apart along the entire dorsal half, with the somata lying near the dorsal surface (Figure 1C). To mimic parameters used in clinical DRGS [4–5], we utilized biphasic stimulation waveforms of a pulse width of 300 µs and a pulse frequency of 40 Hz.

**Figure 4:**
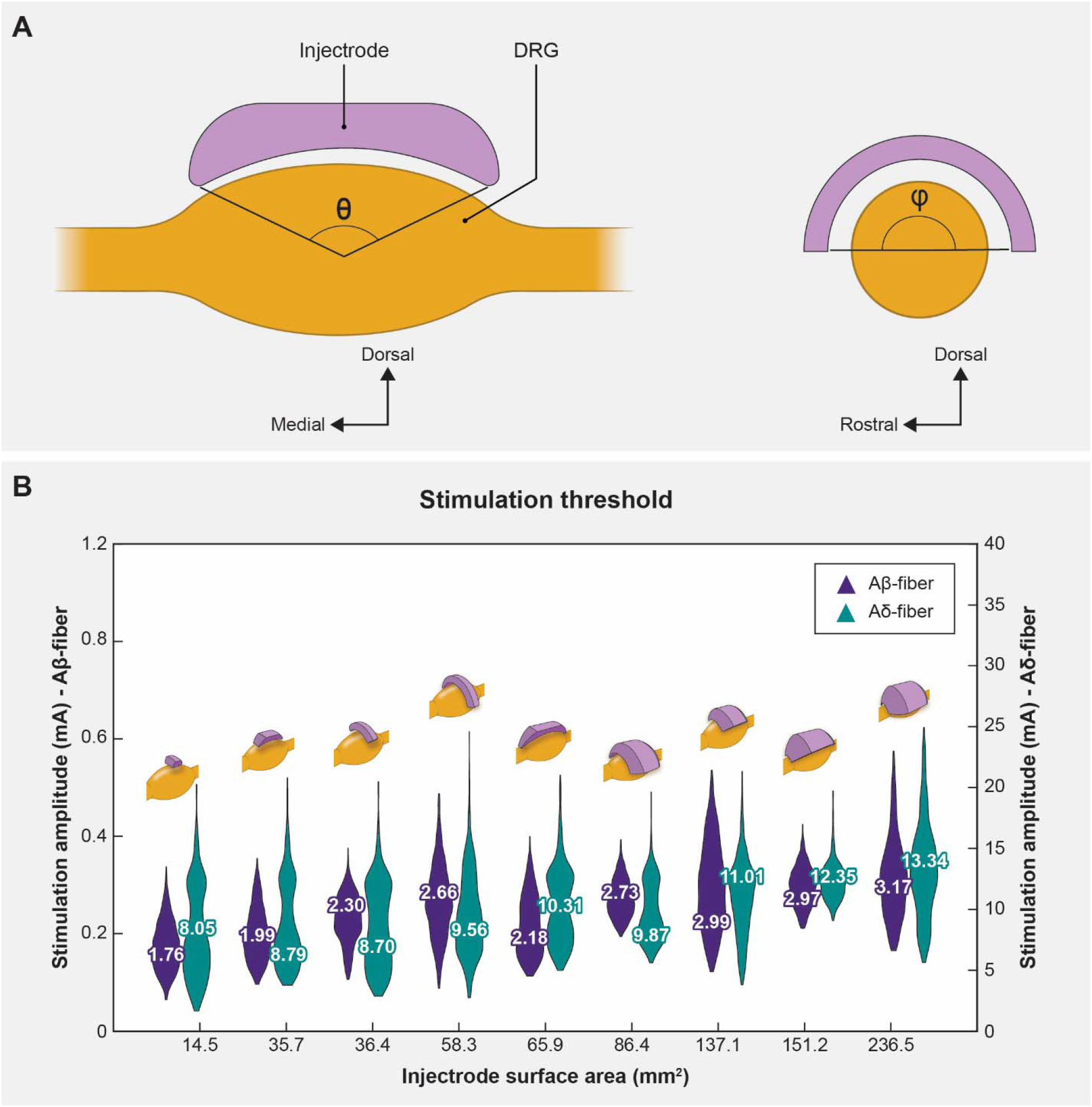
Sagittal and transverse cross sections of the DRG and the Injectrode indicating the various angles of coverage of the Injectrode and the corresponding mean activation thresholds. (A) The angles in both planes (θ,φ) varied from 30 to 150 degrees at an interval of 60 degrees, thus generating a total of nine models. (B) Plots showing comparison between the distribution of activation thresholds of Aβ- and Aδ-fibers generated by the various Injectrode geometries with the mean values inset and the corresponding Injectrode geometry at the top of each violin plot. The left axis refers to the Aβ-fiber amplitudes and the right axis refers to the Aδ-fiber amplitudes, respectively.

**Table 3.** Dimensions of the canonical FEM model of the human L5 DRG.

| Parameter | Value | Reference |
| --- | --- | --- |
| DRG length | 9.40 mm | [40] |
| DRG width | 5.90 mm | [40] |
| Nerve root radius | 1.19 mm | [41] |
| Dural sheath thickness | 0.15 mm | [42] |
| Foramen height | 17.1 mm | [40] |
| Encapsulation layer | 0.30 mm | [30] |

**Table 4.**
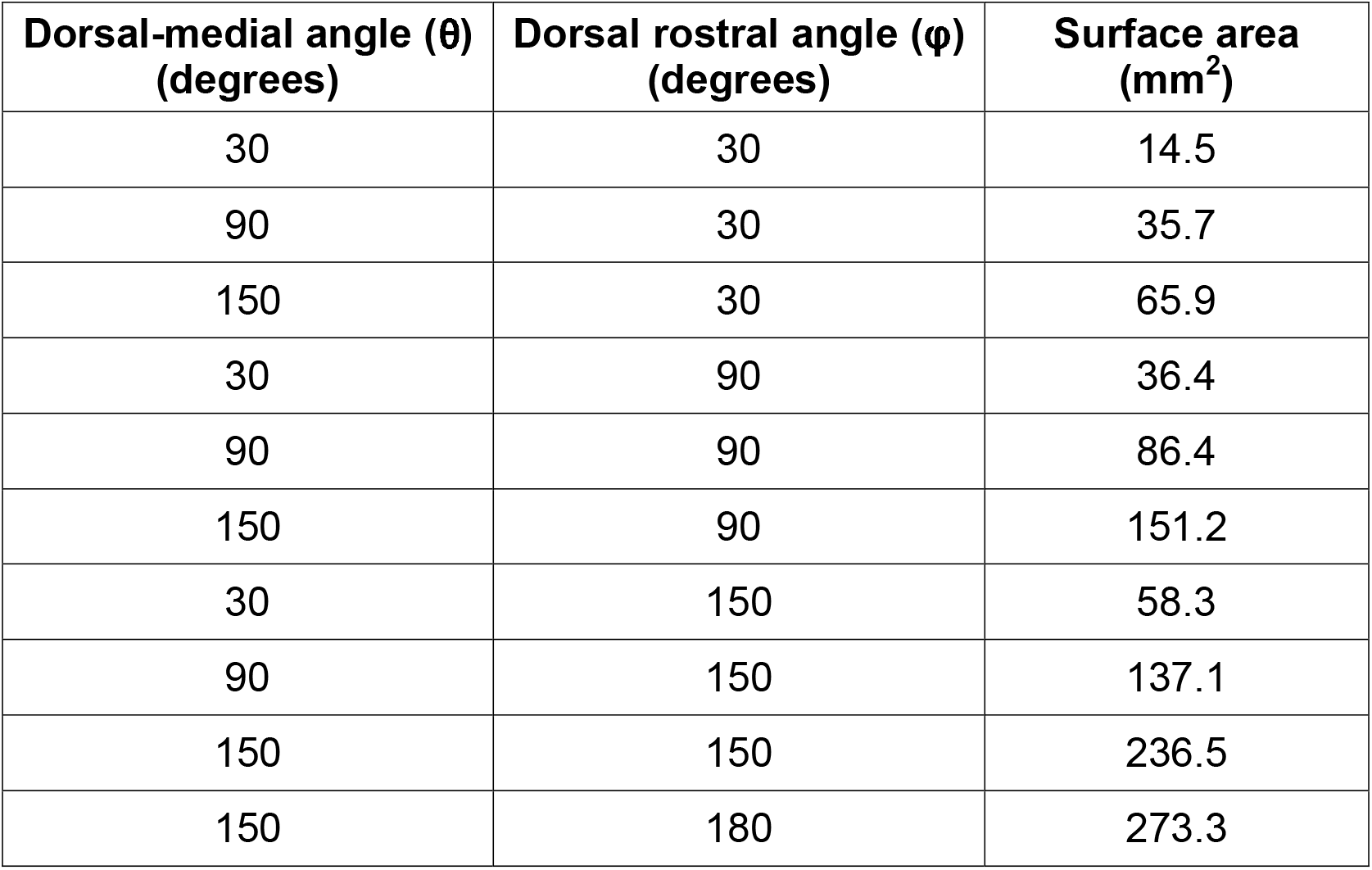
Surface areas of the different Injectrode shapes.

| **Dorsal-medial angle ($\theta$)**<br>**(degrees)** | **Dorsal rostral angle ($\phi$)**<br>**(degrees)** | **Surface area**<br>**(mm<sup>2</sup>)** |
| --- | --- | --- |
| 30 | 30 | 14.5 |
| 90 | 30 | 35.7 |
| 150 | 30 | 65.9 |
| 30 | 90 | 36.4 |
| 90 | 90 | 86.4 |
| 150 | 90 | 151.2 |
| 30 | 150 | 58.3 |
| 90 | 150 | 137.1 |
| 150 | 150 | 236.5 |
| 150 | 180 | 273.3 |

### Model of clinical DRGS applied with a complete Injectrode system

Our final goal was to build a full-scale human model with bilateral lumbar DRG and corresponding spinal nerves to perform model-based design of the Injectrode system (Figure 5). The primary model structures were based on the “Duke” model from the IT’IS foundation virtual population models [37]. We modeled a skin layer with a uniform thickness of 2.64 mm [46] and a fat layer with a uniform thickness of 3.76 mm [45]. For simplicity, we modeled the fat and subcutaneous adipose tissue as a single layer [44]. To represent the rest of the soft tissues in the abdominal region, we assigned them the electrical conductivity of the general thorax [29] (Table 1). Additionally, we included the entire thoracic, lumbar, and sacral vertebrae, along with the spinal cord, dura mater, and cerebrospinal fluid. To alleviate computational demands, our model focused solely on the L5 lumbar roots, as they are a common target for managing chronic foot pain [16–17,25]. The model is comprised of two transcutaneous electrical stimulation (TES) surface electrodes (side length 5 cm) placed equidistant from the central sagittal plane, with a 10 cm edge-to-edge distance, at the L2 vertebral level (Figure 5A). Directly beneath the surface electrodes were circular collectors positioned at a depth of approximately 2 mm from the outer skin [43–44]. These collectors (diameter 5 cm) were connected to the Injectrode that encompassed the L5 DRG via a 0.5 mm diameter wire (Figure 5C). We assigned the collectors, wires, and the Injectrode the electrical conductivity of platinum (Table 1). We insulated the wires and modeled a bipolar Injectrode system with one TES patch modeled as a current stimulus terminal and another set as ground (i.e., 0 V). In our model, a bipolar stimulation configuration involved having two Injectrodes positioned on the DRG on the opposite sides of the same spinal level (L5), each with its corresponding insulated wire, collector, and TES patch. The Injectrode-collector-TES system on the contralateral side acted as a return path. We populated the dorsal half of the ipsilateral L5 DRG near the Injectrode connected to the collector beneath the active terminal with 1378 Aβ-fibers spaced 300 µm apart, with the somata lying near the dorsal surface (Figure 1C). Additionally, to consider the possibility of generating acute pain sensations as a potential side effect of DRGS, we included thinly myelinated medium-diameter Aδ-fibers responsible for both noxious and innocuous sensations. The channel dynamics and morphological structure of these Aδ-fibers were modeled based on our previous work [5]. We included a total of 1378 Aδ-fibers spaced 300 µm apart along the entire dorsal half of the DRG, mirroring the distribution of Aβ-fibers. To mimic previous work with the Injectrode system [43], we determined the activation thresholds for DRG neurons in response to a symmetric biphasic stimulus with a pulse width of 250 µs applied at a pulse frequency of 25 Hz. We simulated DRGS with three Injectrode shapes, where the included angle varied as 30, 90, and 150 degrees in the dorsal-medial plane and 30, 90, and 180 degrees in the dorsal-rostral plane (Table 4). The resultant surface areas of the Injectrode were: 14.5, 86.4, and 273.3 mm^2^, respectively (Figure 6). We also encased the Injectrodes and collectors in a 300 μm thick encapsulation layer [30].

**Figure 5:**
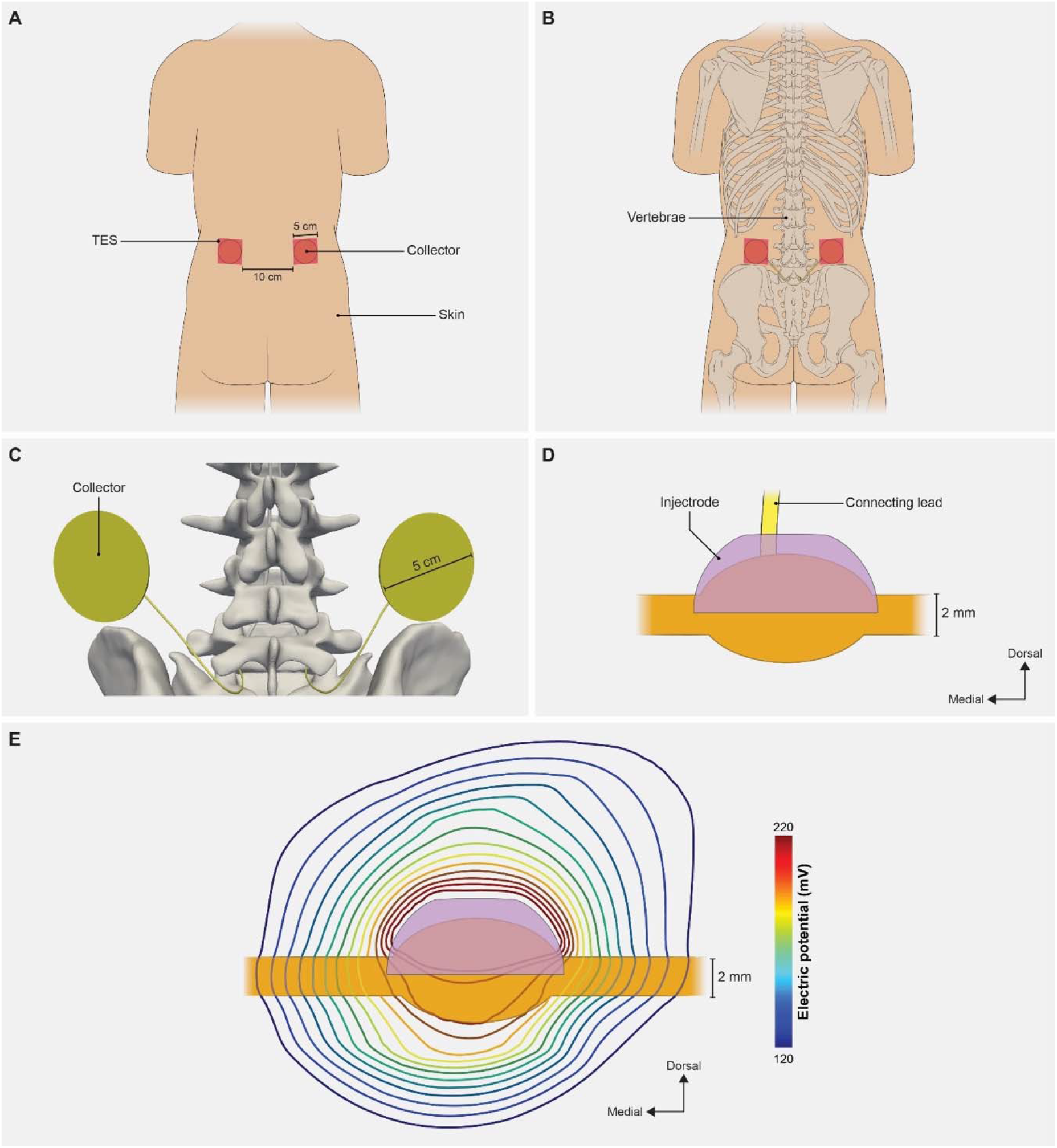
Full-body model with truncated arms, legs and neck with a charge delivery system mimicking clinical implementation of an Injectrode system. (A) Dorsal view of the body with transcutaneous electrical stimulator (TES) patch electrodes visible on the skin surface at the L2 vertebral level. In a bipolar configuration, one TES electrode serves as an active terminal and the other TES electrode is grounded. (B) Collectors are placed directly under the TES patch to receive some of the charge delivered to the TES electrodes. (C) The collectors deliver charge to the Injectrode using a connecting lead made of the same material and inserted in the spinal cavity using an interlaminar approach. (D) A side view of the Injectrode, DRG, and the connecting lead. The Injectrode sits right on top of the dorsal aspect of the DRG. (E) Isopotential lines of the potential field generated by the bipolar TES-collector-Injectrode system near the DRG.

**Figure 6:**
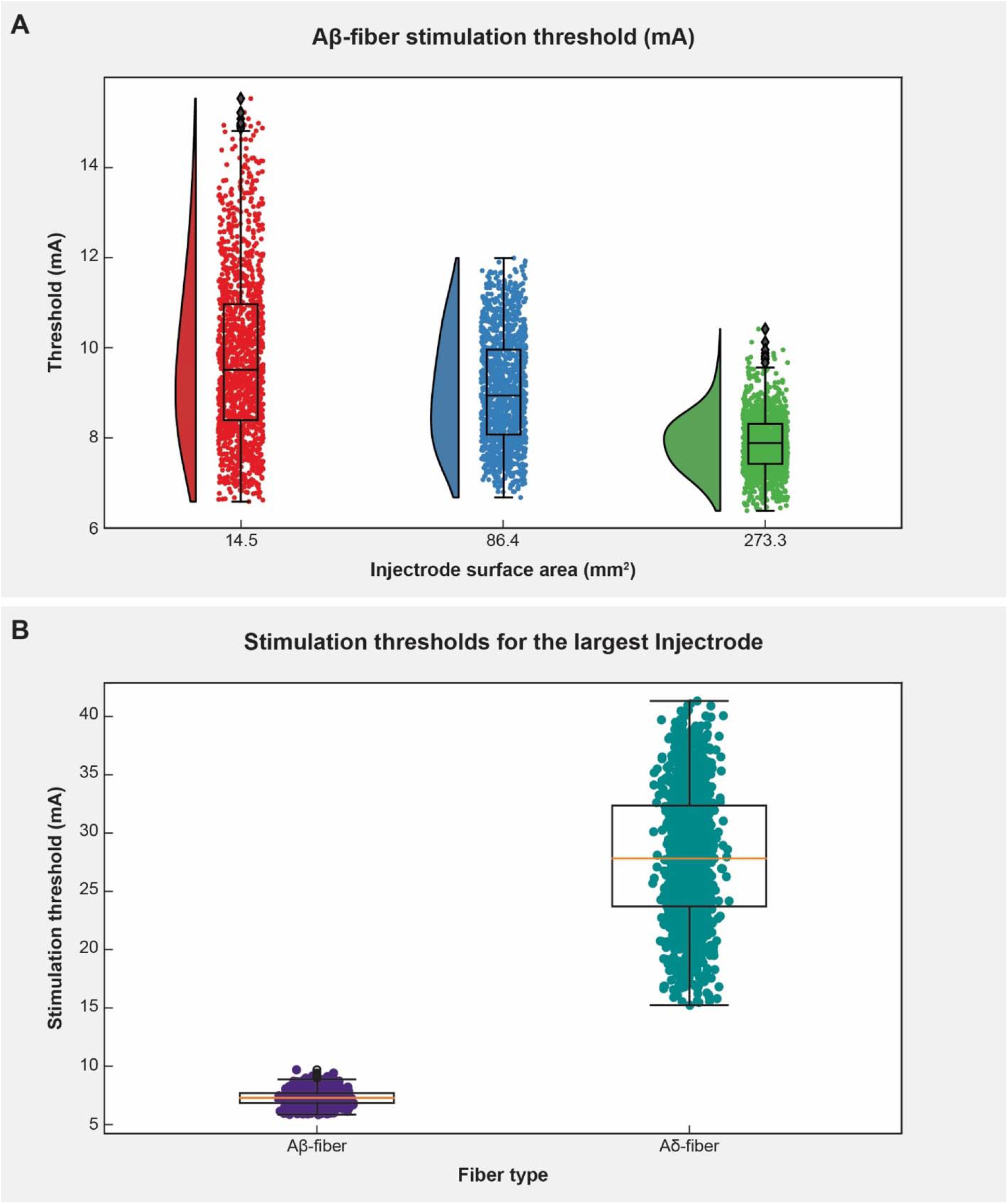
The effect of Injectrode size on activation thresholds with the bipolar Injectrode DRGS configuration in the full-body model. (A) Raincloud plots show the variation of stimulation amplitudes for the entire population of model Aβ-fibers within the DRG. (B) Activation thresholds for the entire populations of Aβ- and Aδ-fibers using the Injectrode with the largest surface area (273.3 mm^2^) considered in this study.

## RESULTS

### Model validation

Stimulus pulse width is a critical parameter when programming a patient’s DRGS system and has been shown to affect neural activation [38] and paresthesia distribution [39] during SCS. Therefore, we scaled the canonical DRG model to match the dimensions of a feline L7 DRG and calculated primary afferent activation thresholds for several pulse widths (i.e., 80, 150, 300 μs) and a constant pulse frequency of 58 Hz [24] with the Injectrode centered above the ganglion in monopolar stimulation conditions. Figure 3 shows the activation threshold as a function of the three pulse widths. The activation thresholds were nearly consistent across the dorsal-rostral plane in a cross-section taken along the width of the DRG, because the Injectrode uniformly covers that area (Figure 3A). However, along the dorsal-medial plane in a cross-section taken along the length of the DRG, the activation thresholds tend to increase with increasing distance of the axons from the Injectrode. This can be attributed to the larger distance between the neuron and the Injectrode (Figure 3A). We also observed the expected decrease in activation thresholds for longer pulse widths (Figure 3B).

To validate our modeling approach, we also compared the activation thresholds predicted with our computational model to the activation thresholds measured in our previous experimental research in a feline model of DRGS with the Injectrode [24], where the L6 and L7 DRG were exposed in four cats via partial laminectomy or burr hole. In this previous work, we stimulated the DRG with an Injectrode using biphasic pulses at three different pulse widths (80, 150, 300 μs) and pulse amplitudes spanning the range used for clinical DRG stimulation. We previously used nerve cuff electrodes to record antidromic evoked compound action potentials (ECAPs) in the sciatic, tibial, and common peroneal nerves. Then we determined the charge-thresholds and recruitment rates for ECAPs from Aα-, Aβ-, and Aδ-fibers. The minimum predicted Aβ-fiber recruitment thresholds from our models have a maximum mean absolute percentage error less than 37.5% of the values measured in these previous acute experiments [24].

### Injectrode geometry

Understanding the impact of changes in surface area on neural recruitment during DRGS is crucial due to the importance of contact surface area as a design parameter, particularly in the case of an Injectrode where the entire surface acts as an active contact and may vary from patient to patient. Therefore, we made nine distinct models with different Injectrode geometries. For monopolar stimulation applied with a stimulus pulse width of 300 µs and pulse frequency of 40 Hz, we observed that increasing the Injectrode surface area almost exclusively increased the mean activation thresholds of Aβ-fibers within the DRG (Figure 4B). The one exception was when the surface area was increased from 58 mm^2^ to 66 mm^2^, during which a minor decrease in mean threshold amplitude was observed. It was also observed that for different models of Injectrode of similar area, the geometries which spanned longer in the dorsal-medial plane (Figure 4B) had relatively lower amplitudes. There was no activation of Aδ-fibers at comparable activation thresholds. In some cases, irrespective of the size of the Injectrode, we observed a small percentage (<10%) of the acute pain fibers activated, when we stimulate the entire population of the Aβ-fiber mechanoreceptors (Figure 4B).

### Effect of clinical parameters on charge delivery via bipolar TES-Collector-Injectrode system

The Injectrode system employs TES electrodes that wirelessly transfer charge to subcutaneous collectors, which is the uncoated wire section of an Injectrode placed under the skin during the final step of the injection procedure. This non-invasive stimulation setup minimizes associated risks and complications inherently part of percutaneous electrode systems [43,53–54]. To investigate the design parameters of the Injectrode for this type of bipolar stimulation, we developed a FEM model of a non-invasive transcutaneous charge delivery system in a full-scale human body model with Injectrodes placed bilaterally at the L5 DRG. We applied stimulation at the TES electrodes, which wirelessly transferred charge to the collectors, which in turn were connected to the Injectrodes via insulated wires. To understand the design parameters affecting the electric field generated by the Injectrode near the DRG, we simulated DRGS with three Injectrode geometries (Table 4; Figure 6A; Figure 7). Our findings demonstrate that increasing the surface area of the Injectrode yields a notable reduction in the mean activation threshold for the Aβ-fiber population within the DRG, as illustrated in Figure 6A. This trend contrasts with the outcomes observed for the canonical models lacking an external TES system. Furthermore, across all three Injectrode models, we observed that none of the Aδ-fibers exhibited any activity until a substantial proportion (>75%) of the entire Aβ-fiber population had been activated (Figure 7). Particularly noteworthy is the Injectrode with the largest surface area, which demonstrated a significant therapeutic window (difference between the minimum activation threshold of Aδ-fibers and the maximum activation threshold of Aβ-fibers) of 7.5 mA compared to the other Injectrodes (Figure 6B, Figure 7C). While a similar trend with a reduced therapeutic window was observed for the medium-sized surface area Injectrode (Figure 7B), the smallest surface area Injectrode exhibited a significant overlap in stimulation amplitude required to activate all Aβ-fibers and a small percentage of Aδ-fibers (Figure 7A).

**Figure 7:**
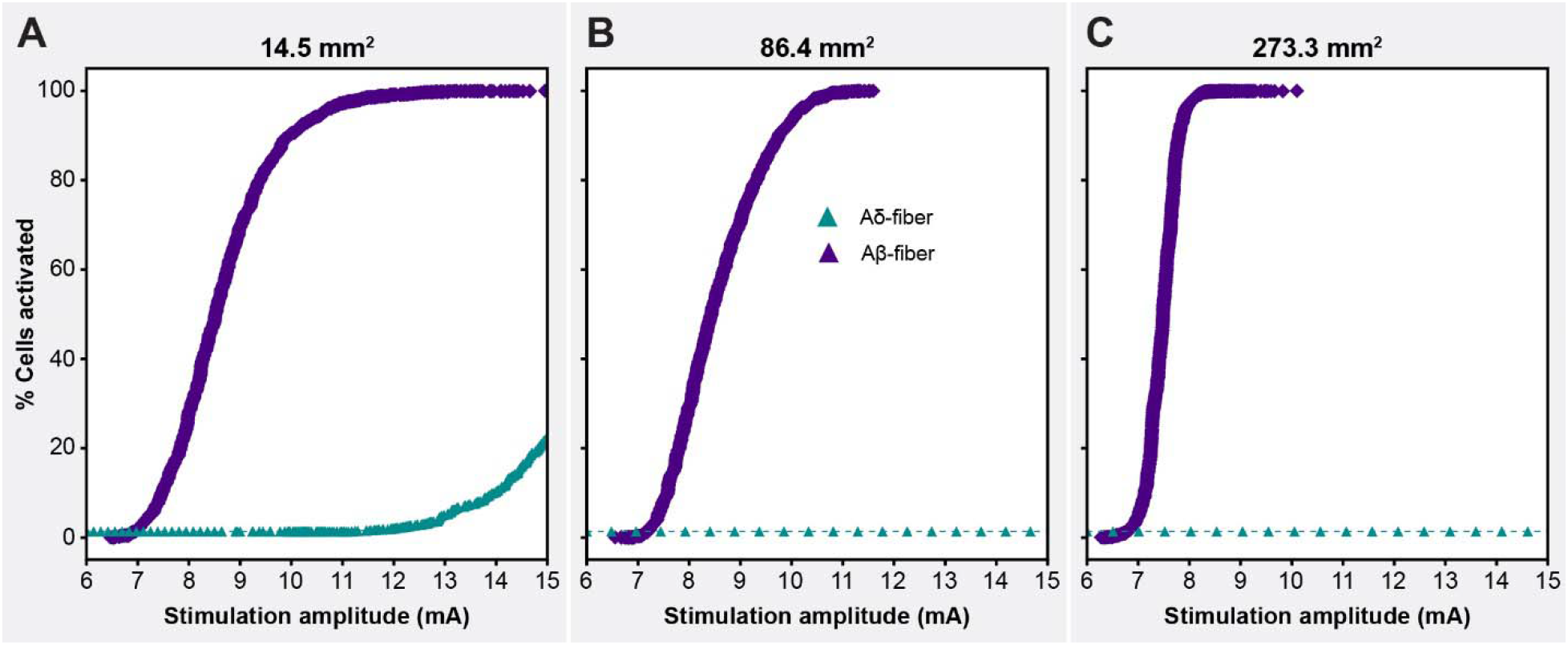
Recruitment curves of the Aβ- and Aδ-fibers for the Injectrodes of three different surface areas; with the maximum amplitude corresponding to the amplitude necessary to activate the entire Aβ-fiber population.

## DISCUSSION

Compared to fully implantable systems, minimally invasive neuromodulation therapies are an advantageous low-cost, low-risk avenue to deliver electrical stimulation to deep neural targets. The Injectrode is composed of a platinum/iridium micro coil, which is designed to be introduced through a needle (18g, 1.27mm) and positioned around a specific neuroanatomical target. Once in place, it assumes a highly conforming and flexible structure, creating a clinical-grade electrode platform [26,43,53]. This unique design is intended to allow the Injectrode to conform around the target neuroanatomy, enabling access to challenging anatomical sites, including the DRG. Prior investigations conducted by our group have consistently demonstrated that clinical DRGS primarily impacts the functioning of large-diameter myelinated Aβ-fibers, without directly activating medium-diameter thinly myelinated Aδ-fibers responsible for noxious or innocuous sensation or small-diameter unmyelinated C-fibers associated with pain perception [3–5]. These findings align with experimental studies which have further substantiated that DRGS activates neurons exhibiting conduction velocities primarily akin to Aβ-fibers [27]. Therefore, since DRGS is believed to provide analgesia via pain-gating induced by the activation of Aβ-fibers [3–5], we built computational models to study neural recruitment of Aβ-fibers within the DRG using the Injectrode.

While previous studies have investigated the use of the Injectrode for vagus nerve stimulation in preclinical models [43] or DRGS using clinical leads [3–4], the specific impact of clinically adjustable parameters on neural recruitment during DRGS with the Injectrode system has yet to be fully understood. The DRG, located in the spinal foramen, is a much deeper neural target compared to the vagus nerve [43]. Computational models of SCS [51] demonstrated that when electrodes are placed directly under the vertebrae, the current distribution is directed away from the highly resistive vertebral bones and towards the neural target. Therefore, the Injectrode placed within the partially enclosed and highly resistive compartmental structure of the foramen, will exhibit distinct neural activity patterns. Additionally, the anatomical organization of the DRG and the vagus nerve differ significantly. In the DRG, sensory axons and somata are positioned dorsally and are completely segregated from motor axons [50]. Conversely, the vagus nerve contains both sensory and motor fibers, with sensory axons predominantly located in the dorsal aspect of the nerve, while motor and sympathetic efferents are found ventrally, playing a role in organ control and autonomic responses. In our previous modeling efforts, the vagus nerve is modeled with straight axons [43]. In contrast, the DRG neurons are pseudounipolar, characterized by a single axon process that extends from the soma, bifurcates at a large node of Ranvier called the T-junction, and gives rise to an axon projecting to the spinal cord and another axon extending to the periphery [11], resulting in less predictable activation patterns. To address this knowledge gap, we leveraged computational modeling to optimize the clinical efficacy of the system and explore various electrode configurations. Since the Injectrode differs from a standard clinical electrode through its flexible shape and orientation, it is important to test neural activation outcomes considering different geometries of the Injectrode itself. The results presented here serve as a crucial step towards enhancing our understanding of the direct neural response to DRGS with the Injectrode and ultimately improving patient outcomes.

### Model validation

Our first step was to validate our computational modeling approach by comparing our model predictions to our previous experimental work of DRGS using the Injectrode [24]. The contour maps of stimulation amplitude clearly illustrate that the threshold values remain relatively consistent across the dorsal-rostral cross-section taken along the center of the DRG, while exhibiting variation along the dorsal-medial plane (Figure 3). This notable difference can be attributed to the unique shape of the Injectrode, which was designed based on the surface area utilized in the acute experiments [24]. As a result, the Injectrode covers the region of the DRG containing all the cell bodies and bifurcations of the axons within the dorsal-rostral cross-section, exerting direct influence over them due to its nearly complete width coverage of the DRG (Figure 3A). In contrast, the length of the Injectrode is comparatively shorter than the length of the DRG, resulting in numerous axons having their somata positioned beyond the region of direct influence exerted by the Injectrode (Figure 3A). This distinction clarifies the observed minor variations in threshold values and emphasizes the importance of considering the spatial dimensions of the Injectrode when assessing its impact on neuronal activation. As expected, the minimum predicted Aβ-fiber recruitment thresholds from our model decreased with increasing pulse widths (Figure 3B). In comparison to data from the acute experiments in which antidromic ECAPs were recorded, we observed a mean absolute percentage error ranging from a minimum of 5.2% to a maximum of 37.5% across the three tested pulse widths. The findings of this study indicate that, despite the limitations of an open cut-down procedure as posed by fluctuating electrical conductivities from fluid build-up and bleeding in the acute experiments, the computational model successfully replicated the experimental results with a substantial level of concordance. The computational models presented in this work can thus serve as a robust platform to investigate various parameters and configurations essential for the effective clinical deployment of the Injectrode.

### The influence of monopolar Injectrode size on DRGS

After validating our model, we then proceeded to build a computational model of a canonical human L5 DRG with nine different Injectrode geometries (Figure 4). Our analysis reaffirmed the commonly understood principle that an increase in surface area is accompanied by a decrease in charge density. Consequently, in the case of Injectrode geometries of larger surface area, a higher stimulation amplitude was required to effectively activate the Aβ-fibers. However, an interesting exception emerged when the surface area was expanded from 58 mm^2^ to 66 mm^2^, resulting in a minor but substantial reduction in the mean threshold amplitude. This exception can be attributed to the fact that the Injectrode with a surface area of 66 mm^2^ spans a longer distance in the dorsal-medial plane (Figure 4), thereby exerting direct influence over a greater number of fibers. This observation is supported by a consistent trend in our results, where Injectrode geometries with similar surface areas exhibited relatively lower amplitudes when they spanned longer distances in the dorsal-medial plane (Figure 4). This trend can be explained by the fact that the length of the dorsal root ganglia (DRG) is greater than its width (Table 3). As a result, when the Injectrode geometries are centrally positioned on the dorsal aspect of the DRG, those geometries spanning along the length of the DRG will invariably impact more fibers compared to those spanning along the width.

Our study provides valuable insights into the activation profiles of Aδ- and Aβ-fibers using the Injectrode in a clinical context. We observed that the activation thresholds for Aδ-fibers were generally higher compared to the majority of Aβ-fibers. Consequently, the therapeutic window for the Injectrode is considerable, as no significant activation of Aδ-fibers occurred at activation thresholds comparable to most Aβ-fibers. It is worth noting that in some cases, a small fraction (<10%) of Aδ-fibers was activated when all Aβ-fibers were stimulated. However, we believe that this occurrence is unlikely to be clinically relevant. Only in the scenario involving the smallest Injectrode size did we encounter an instance where more than 5% of Aδ-fibers were activated alongside all Aβ-fibers. To mitigate this issue in clinical applications, we propose two potential solutions. Firstly, utilizing a larger Injectrode size, as there was little-to-no Aδ-fiber activity observed for stimulation amplitudes required to activate the entire Aβ-fiber population from Injectrodes with larger electrode surface areas. Secondly, stimulating a portion of Aβ-fiber population would be more clinically relevant since it is not necessary to activate all the Aβ-fibers to produce analgesia for pain relief [4,5]. It is well demonstrated in our results that no Injectrode model activated any Aδ-fibers before at least 75% of the Aβ-fibers were activated. Moreover, the charge densities for the largest and smallest Injectrode were calculated to be 0.40 μC/cm² and 3.64 μC/cm², respectively. It is worth noting that both charge densities are substantially lower than the charge densities required for activation with clinical DRGS electrodes that we estimated in previous computational modeling work (10 μC/cm²) [4,5], suggesting a larger therapeutic window before activation of off-target fibers, as demonstrated by our results. In a clinical scenario, having an electrode with lower charge density is crucial to minimize tissue damage at the electrode-tissue interface by reducing the likelihood of electrochemical reactions, electrolysis, and pH changes. This promotes the safety and long-term viability of the electrode. Moreover, lower charge density enables improved selectivity in neural stimulation, allowing for precise control over the activation threshold of specific neural populations and improved battery life if connected to an implantable pulse generator (IPG). This targeted stimulation minimizes unintended activation of nearby or non-targeted neural structures, with the potential to lead to more accurate and effective neuromodulation.

### Injectrode size changes transcutaneous impedance pathways, altering shunting between TES patches

Although results from the canonical model suggested that using a large surface area Injectrode during monopolar stimulation would lower the charge density necessary for neural activation and subsequently provide an increased therapeutic window and reduce the risk of tissue damage, it was still unclear how the use of the Injectrode in a bipolar DRGS configuration would affect charge delivery in a clinical setting. Both experimental and previous modeling studies have demonstrated that the placement of a second Injectrode on a neural target significantly increases the proportion of the applied electric field that reaches the neural target [43]. This improvement can be attributed to the impedance characteristics of the pathways between the active Injectrode, the “return electrode,” and the impedance between two TES patches overlaying the two collectors. Experimental results indicate that the deep penetration of current in a monopolar configuration (single Injectrode) is approximately 5%, whereas in a bipolar configuration (two Injectrodes), it increases to 20% [43,53].

The Injectrode system allows for the injection of the second “return” Injectrode at any desired location and size, and the ease of injection and removal further enhances its utility [55]. This feature enables the possibility of redirecting current to specific areas as needed. Moreover, the Injectrode involves the TES electrodes wirelessly transferring charge to the subcutaneous collectors which in turn are connected to the Injectrodes via insulated wires, which are coated portions of the same lead. This non-invasive system reduces the potential for battery and device-related complications, including recalls and the need for surgical removal. Thus, non-invasive stimulation methods are thought to provide a safer, more accessible, and reversible approach, minimizing the risk of infections, and simplifying management.

In our model, the bipolar configuration was defined as having another Injectrode at the contralateral DRG, which consisted of its corresponding set of electrode, coated wire portion, collector, and TES patch (Figure 5C). This entire system acted as a return path. It is imperative to mention that in the full body Injectrode-collector-TES patch set-up, a monopolar configuration can be defined as having only one Injectrode on the target DRG, and the ‘return’ TES patch having no associated Injectrode/collector (not examined in this study). In what we define as bipolar configuration (as discussed in Section 2.3 and Section 3.3), we had two Injectrodes near the two DRG at the same level (L5) and both TES patches have associated collectors which connect to these two Injectrodes via insulated wires (Figure 5D).

We observed that in the case of charge delivery using the entire TES patch-collector-lead-electrode Injectrode system, an increase in size of both the active and return Injectrode led to a decrease in activation thresholds (Figure 6A). The activation threshold plots reveal that when using an Injectrode with a larger surface area, both the average thresholds and the variance of thresholds are significantly reduced (Figure 6A). This indicates that the larger Injectrode encompasses a greater number of axons within its direct influence, highlighting an important aspect of design consideration. This finding also contradicted what we had previously observed in our monopolar simulations. This discrepancy can be attributed to the manner in which stimulation occurs. In the monopolar stimulation setup, the stimulus current is directly applied to the surface of the Injectrode (Figures 1 and 3) and the size of Injectrode (and therefore impedance) does not change the ‘return path’ for the active electrode to the ground electrode which in this case is the grounded thorax. However, in the bipolar stimulation setup, the current is applied to the TES patch, and it wirelessly traverses to the subcutaneous collector. Subsequently, the collector is connected to the Injectrode via coated and connected lead portion of the device (Figure 5). For an Injectrode with a smaller surface area, there is an increase in electrode impedance, which ultimately results in less current traveling to the smaller Injectrode. Thus, the electric field reaching the DRG also becomes weaker, resulting in higher activation thresholds. Similarly, an increase in the size of the Injectrode lowers the impedance path, causing more current to go deep than shunting between TES patches. However, this result also highlights a potential limitation of isolated-component canonical models. Consequently, it emphasizes the need for a comprehensive full-body model that encompasses the complete charge delivery system.

It was also observed that the two Injectrodes implanted near the two DRG are not necessarily near enough to each other to impact the spread of the electric field from the working electrode. This differs from the conventional definition of bipolar configuration used for multi-contact SCS or DRG leads [4–5], where the working and reference electrodes/contacts are placed close enough such that the spread of the field from the working electrode is significantly impacted. In this scenario, significant current shunts between the working electrode and reference electrode instead of spreading out uniformly from the working electrode. This is why the fall off of an electric field is 1/r in a traditional monopolar configuration, where r is the distance from the working electrode, whereas it is 1/r^2^ in a traditional bipolar configuration. The spacing of the bipolar Injectrodes modeled in this study is such that the field potentials generated by the working electrode at the DRG is effectively monopolar with an electric field fall off of 1/r. However, the reason for employing a second Injectrode is rooted in the utilization of a biphasic waveform. This approach allows us to achieve Aβ-fiber activation at two dorsal root ganglia (DRGs) instead of just one. This could prove beneficial, especially in cases of widespread pain or bilateral pain within a clinical context. Injecting into two DRGs becomes more viable due to the simplicity of the injection process, as compared to the placement of two conventional multi-contact leads targeting multiple DRGs. Alternatively, for upcoming applications, positioning the return Injectrode outside the neural foramen is also a possibility. This approach would offer the advantage of creating a larger return electrode, thereby reducing unintended activation through decreased charge density. Moreover, it can be designed to enhance the efficiency of transferring transcutaneous electrical stimulation to the deeper target Injectrode by decreasing impedance in the deep return pathway.

### Reduced charge density of Injectrode compared to clinical leads

In comparison to the stimulus amplitudes required for response using standard clinical leads implanted in the intraforaminal space [4–5], the Injectrode with TES patches in our study necessitated considerably higher amplitudes. However, it is important to note that the charge density associated with the Injectrode varies significantly depending on the surface area. Specifically, the Injectrode with the largest surface area exhibited a charge density of 0.705 μC/cm^2^, while the smallest surface area resulted in a higher charge density of 16.3 μC/cm^2^. In contrast, the charge density associated with clinical leads was estimated to be 10 μC/cm^2^ in previous computational modeling work [4–5]. These findings indicate that the Injectrode with the largest surface area provides a lower charge density, potentially offering a larger therapeutic window and reducing the likelihood of off-target fiber activation and the risk of tissue damage. Moreover, the model-based activation thresholds are within the range of amplitudes described in existing literature required to elicit a response in cases where similar surface electrodes were used, e.g., vagus nerve stimulation [43].

An essential pathway for side effects in DRGS is the potential activation of Aδ- and C-fibers, which can lead to undesirable side effects. Previous modeling studies have indicated that unmyelinated C-fibers possess higher activation thresholds and are not activated during DRGS [4]; however, there may still be situations where Aδ-fibers are activated, particularly with longer pulse width stimulation at clinical amplitudes [5]. In our current model, we observe that Aδ-fibers are not activated until a significant percentage (>75%) of myelinated large-diameter Aβ-fibers have been activated (Figure 7). It is important to note that only in the case of the smallest Injectrode, there was an overlap between stimulation amplitudes needed to activate the entire population of Aβ-fibers and the activation of any Aδ-fibers (Figure 7A). There is a considerable therapeutic window for the larger surface area Injectrodes where the entire Aβ-fiber population can be activated before activating any Aδ-fibers (Figure 7B, Figure 7C). This finding underscores the importance of ensuring that the electrode encompasses as much surface area of the DRG as possible to optimize the therapeutic effects of stimulation. A large therapeutic window between Aβ- and Aδ-fiber activation can be highly advantageous in clinical applications, minimizing the risk of unwanted nociceptive responses and enhancing the overall effectiveness and tolerability of DRGS treatment.

### Limitations

It is also important to acknowledge the potential limitations of our computational modeling approach. While we utilized previously published clinical and experimental data, there are several considerations to keep in mind. Firstly, our canonical models represented anatomical compartments, such as foraminal bone and intraforaminal tissue, as simplified concentric cylinders around the feline/human DRG. Although this approach has been commonly used in studying other neurostimulation therapies, our full-body model demonstrated that the complex anatomy of soft tissues and the path of charge delivery can impact the predictions of computational models of DRGS. Future studies could benefit from employing a patient-specific modeling approach [49], to better understand how the intricate anatomy of the spinal column affects DRGS model predictions. Our assumption of an idealized trajectory for axons within the ganglion might not fully capture the reality of stem axons, which are complex and winding, forming tightly packed glomeruli around somata before bifurcating into central and peripheral axons [11]. The influence of tightly coiled stem axons on DRGS thresholds remains unknown. Further research should explore the effects of intricate stem axon trajectories and the functional organization of cells within the human DRG on neuronal activation, and how these factors impact DRGS. Lastly, it’s important to note that the charge delivery system we modeled, consisting of the TES patch, collector, lead, and electrode of the Injectrode, is only a representative model in our simulations. In particular, the deployment process of the Injectrode is intricate, and its shape and volume are determined as a result of this process. While the ideal scenario would be to have an Injectrode that fully encompasses the dorsal aspect of the DRG, practical implementation procedures may impose limitations. In forthcoming computational and functional investigations, enhancing the fidelity of our models could involve the integration of CT scan-based representations of previously deployed Injectrode configurations. This refinement would facilitate the faithful recreation of viable shapes and offer an avenue for exploring the phenomenon of soft tissue scarring surrounding the Injectrode and the collector. Additionally, it’s crucial to recognize that expanding the dimensions of the Injectrode to envelop the dorsal aspect of the DRG warrants careful consideration. This adjustment may potentially result in the proximity of the active electrode to ventral motor efferents, raising potential implications for the activation of motor fibers. This approach would provide a more realistic representation of the actual configuration of the Injectrode and improve the accuracy of the simulations.

## CONCLUSION

Our modeling shows that the Injectrode is a clinically viable technology for minimally invasive stimulation of deep neural targets, such as the DRG. A wireless sub-cutaneous collector-based charge delivery system is able to recruit DRG neurons. Further, based on our findings, the orientation and size of the Injectrode are crucial factors in effectively stimulating deep neural targets. Larger Injectrodes provide an extended therapeutic window prior to unintentional fiber activation, while smaller Injectrodes behave akin to conventional DRG electrodes resembling a point-source electrode. These results underscore the importance of thoughtful electrode placement in achieving optimal outcomes in DRGS therapy. In light of the promising results of our study, future research efforts could benefit from a more comprehensive exploration of the different parameters involved, using a design of experiments approach, to fully explore the potential of DRGS as a treatment for chronic pain.

## ACKNOWLEGDEMENTS

This work was supported by a grant from the National Institute of Biomedical Imaging and Bioengineering [U18 EB029251] as part of the National Institutes of Health Helping to End Addiction Long-term (HEAL) Initiative, and computational resources were provided by Advanced Research Computing (ARC), a division of Information and Technology Services (ITS) at the University of Michigan, Ann Arbor.

